# Correspondence-aware manifold learning for microscopic and spatial omics imaging: a novel data fusion method bringing MSI to a cellular resolution

**DOI:** 10.1101/2020.09.28.317073

**Authors:** Tina Smets, Tom De Keyser, Thomas Tousseyn, Etienne Waelkens, Bart De Moor

**Affiliations:** STADIUS Center for Dynamical Systems, Signal Processing, and Data Analytics, Department of Electrical Engineering (ESAT), KU Leuven, 3001 Leuven, Belgium; Independent Researcher, 3010 Leuven, Belgium; Department of Pathology, University Hospitals KU Leuven, 3000 Leuven, Belgium; Department of Cellular and Molecular Medicine, KU Leuven, 3000 Leuven, Belgium

## Abstract

High-dimensional molecular measurements are transforming the field of pathology into a data-driven discipline. While H&E stainings are still the gold standard to diagnose disease, the integration of microscopic and molecular information is becoming crucial to advance our understanding of tissue heterogeneity. To this end, we propose a data fusion method that integrates spatial omics and microscopic data obtained from the same tissue slide. Through correspondence-aware manifold learning, we can visualise the biological trends observed in the high-dimensional omics data at microscopic resolution. While data fusion enables the detection of elements that would not be detected taking into account the separate data modalities individually, out-of-sample prediction makes it possible to predict molecular trends outside of the measured tissue area. The proposed dimensionality reduction-based data fusion paradigm will therefore be helpful in deciphering molecular heterogeneity by bringing molecular measurements such as MSI to the cellular resolution.

## 1 Introduction

Pathologists have been relying on morphology-based methods for decades to study and diagnose disease. While such staining approaches enable the assessment of one or two markers in a single tissue slide, spatial transcriptomic and proteomic studies make it possible to evaluate many thousands of molecules simultaneously. The number of studies gathering high-dimensional omics measurements keeps growing in an effort to understand the complex interactions taking place in biological systems. These studies have moved from focusing on single components (e.g. gene) to encompassing the entire genome, and even evaluating complementary omics measurements in parallel (e.g. transcriptomics, proteomics,..) [1, 2].

Increasingly, these components are being evaluated in terms of their spatial organisation as well. A prominent example is the field of Mass Spectrometry Imaging (MSI), which is capable to detect thousands of endogenous (small metabolites, lipids, peptides and proteins) or exogenous (drugs and drug metabolites) species in their spatial context [3]. Another important example is the spatial transcriptomics field which spatially resolves the distribution of gene expression profiles to improve our molecular understanding of tissues [4].

Often, alongside these molecular measurements other imaging modalities, such as high-resolution microscopy images, are being collected as well. Different modalities obtained from the same sample can provide relevant complementary information that cannot be obtained from a single modality [5]. Typically, histological or microscopic images (eg. haematoxylin and eosin stainings) are overlaid with the molecular profiles obtained by MSI. These microscopy images give insight into the correlation between structure and (pathological) function of cells and tissues that is complementary to the molecular information obtained using molecular imaging. Creating these overlays however is challenging because it is difficult to register two images obtained at different spatial resolutions. It is therefore important to overcome these challenges and truly integrate these heterogeneous data sources, a concept referred to as data fusion. By generating a single view or image from a set of source images, we can get a more complete picture of the complex interdependencies present in biological phenomena. In this paper we present a novel data fusion method called ‘correspondence-aware manifold learning’ that builds on recent developments in the dimensionality reduction field.

Non-linear dimensionality reduction methods such as t-distributed Stochastic Neighbor Embedding (t-SNE) [6] are often used for the visualisation of high-dimensional biological data [7]. Not only are these methods capable of detecting non-linear trends, they can also capture the complete feature space when reducing data to two or three components, which is not always the case for methods such as for example Principal Component Analysis (PCA) [8]. Recently, Uniform Manifold Approximation and Projection (UMAP) [9] was introduced to this family of methods with major improvements in terms of scalability, enabling the analysis of large spatial omics data such as MSI.

In earlier work, we have shown how the hyperspectral visualisations obtained using UMAP can reflect the molecular trends present in an entire tissue sample [10]. Connecting these molecular trends to histological information is essential to support clinical settings. These images, obtained from the same subject or tissue sample but acquired in different ways, are expected to show some level of correspondence. The same anatomical structures can be displayed in the microscopic image but can also be reflected by the molecular trends. Where the microscopic information will typically have a lower ‘chemical resolution’ but a very high spatial resolution, the molecular trends are complementary in this regard, as they offer very rich chemical information but at a lower spatial resolution. With correspondence-aware manifold learning we are now able to visualise these molecular trends at a higher spatial resolution by fusing the molecular and microscopic data (Figure 1). Moreover, using out-of-sample prediction we can predict the distribution of these molecules for regions not measured by the molecular measurements such that we can enrich a complete microscopy slide with the biological signals available. We demonstrate our approach for the fusion of representative spatial omics and optical microscopy data.

**Figure 1:**
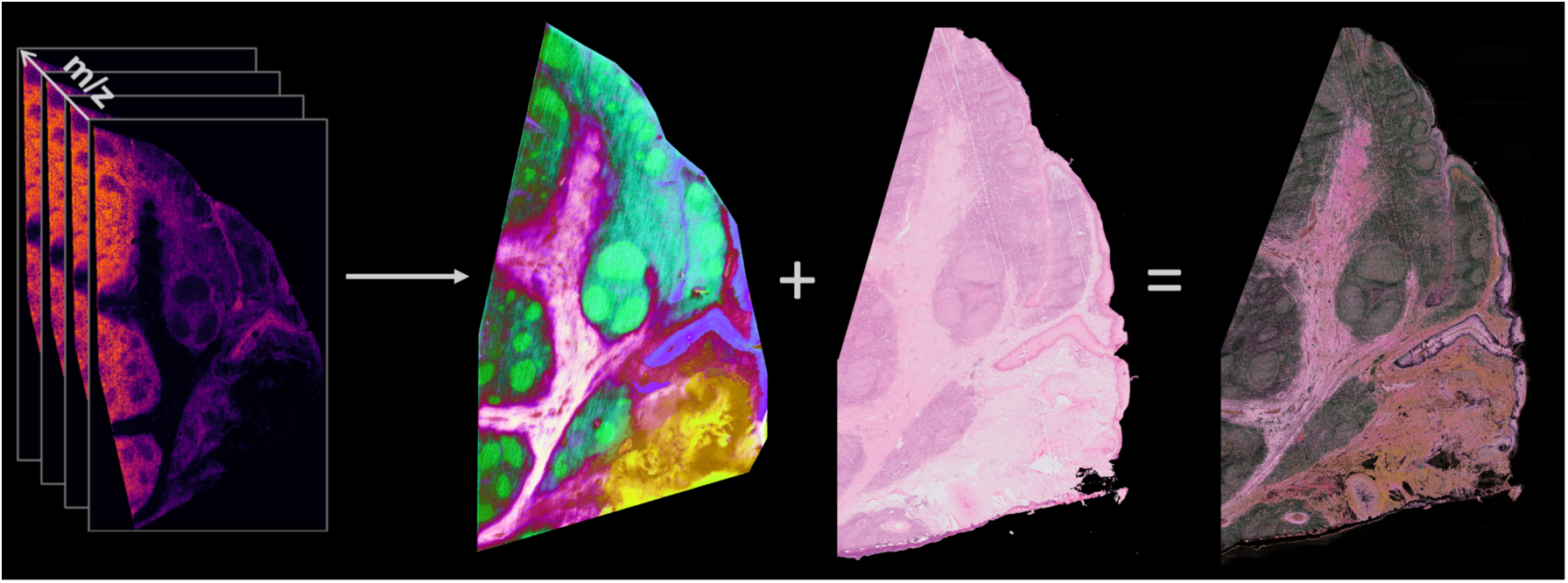
Conceptual overview. Correspondence-aware manifold learning is able to fuse the complete molecular feature space of high-dimensional omics data with their corresponding microscopy image. An example is shown for MSI data where the dataset is first reduced to three dimensions and this representation is fused with the microscopy data such that all molecular trends are visualized at a much higher resolution.

## 2 Methods

### 2.1 UMAP outline

UMAP creates a topological structure that represents the high-dimensional data by assembling approximations of local manifolds, and assembles an equivalent topological structure for a low-dimensional representation of the data. It then optimizes the low-dimensional representation to the high-dimensional data by minimizing the cross entropy between the two topological structures [9]. The algorithm innovates by its mathematical foundations that allows it to make interesting assumptions on the data. An important assumption often used in manifold approximation is a uniform distribution of the data on the manifold [11]. For real world data this is usually not the case. UMAP addresses this problem by creating local Riemannian manifold approximations on which the data is assumed to be uniformly distributed and patching them together into a fuzzy simplicial set representation of the data.

UMAP uses fuzzy set cross entropy to compare the two fuzzy simplicial set representations, (*A, v*) for the high-dimensional data and (*A, w*) for the low-dimensional data. A low-dimensional embedding can be optimized using the cross entropy loss with the membership strength functions v and w as catalysts for attraction and repulsion:

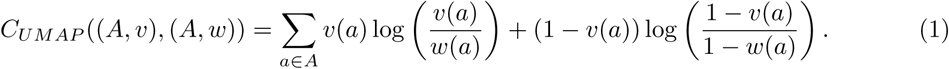

In general UMAP fits well within the family of algorithms such as t-SNE [6] or LargeVis [12]. These algorithms rely on different mathematical principles although their implementations have lots of common ground. Like t-SNE and LargeVis, manifold approximations are implemented as weighted k-neighbour graphs. It is explained in detail in [9] that the main equations from these algorithms also share similarities. Any of these algorithms would suit the data fusion method explained here.

In earlier work we have shown the strong visualisation capabilities of UMAP for MSI data, making it an excellent choice as a general purpose algorithm for high-dimensional omics data [10]. UMAP is therefore used and extended to fit the desired data fusion goals. We use UMAP as a dimensionality reduction algorithm for high-dimensional omics data and adapt UMAP to fuse the resulting low-dimensional representation with high-resolution imaging.

### 2.2 Capturing correspondence

Our goal is to capture spatial correspondence between high-resolution and high-dimensional data and leverage this information into the manifold learning process. Let us use the matrix *A*_*n*×*p*_ to denote the flattened high-resolution data and *B*_*m*×*q*_ for the low-resolution spectral data. The correspondence between these two data sets is recorded in the matrix *C*_*n*×*m*_ such that

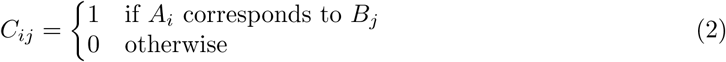

Finding matching pairs can be achieved with registration techniques. Geometric transformation algorithms can estimate a projection between the two coordinate spaces using only a small set of matching pixel pairs. The output is a transformation matrix that can be applied to warp all other pixels from one image to the other.

### 2.3 Correspondence-aware manifold learning (CAML)

Correspondence-aware manifold learning (CAML) creates a fused representation of two datasets such that its corresponding instances lie close to each other in the fused representation. Specifically, CAML models the fusion task as an optimisation problem that aims to (1) preserve local distances within the first dataset, and to (2) minimize distances of corresponding instances with the second dataset.

Consider the task of fusing high-resolution data matrix *A*_*n*×*p*_ and high-dimensional data matrix *B*_*m*×*q*_ based on a correspondence matrix *C*_*n*×*m*_. The information of the correspondence matrix *C*_*n*×*m*_ is reconstructed as a mapping *γ* between the index sets of both datasets.

#### Definition 1

*Define γ* : *I* → 2^*J*^, *the correspondence map for matrices A*_*n*×*p*_ *and B*_*m*×*q*_ *and their respective index sets I* = {1, 2,…, *n*} *and J* = {1, 2,…, *m*}, *such that*

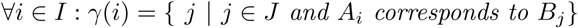

*As a consequence, γ* (*i*) = *ϕ when there are no corresponding instances for A*_*i*_ *in B*.

We capture the concept of distance between corresponding points as an interplay of attraction and repulsion. CAML aims to minimize the repulsion between corresponding points. We formally define the repulsion between *A* and *B*:

#### Definition 2

*Consider matrices A*_*n*×*d*_ *and B*_*m*×*d*_. *Let γ be the correspondence map between the index sets of A and B. Define Ψ*: ℝ^*d*^ → ℝ, *the repulsive strength between these matrices, as*

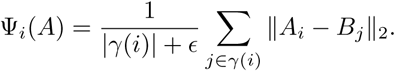

The repulsion *Ψ* is an average because a single instance in *A*_*n*×*p*_ can have multiple corresponding instances in *B*_*m*×*q*_ when *n*< *m*. All corresponding instances receive equal weight. A small number *ϵ* is used to avoid division by zero when calculating the average.

Definition 2 defines repulsion by comparing instances of the two datasets, which is done by choosing a shared dimension *d* for both datasets. In our case, *d* = *p* = 3, similar to RGB color space. The second dataset can then be reduced to *d* dimensions using any dimensionality reduction algorithm, resulting into the two datasets two datasets *A*_*n*×*d*_ and *B*_*m*×*d*_.

Given two datasets *A*_*n*×*d*_ and *B*_*m*×*d*_ and their correspondence map *γ*, CAML can be formulated as a constrained optimization problem for a manifold learning cost function *C*_*MAN*_.

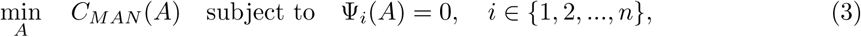

This equality-constrained problem can be transformed into the following quadratic penalty function, which concludes the CAML cost function:

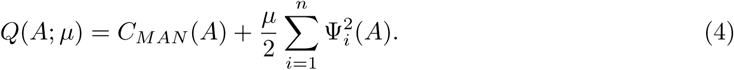

The first term in Equation 4 focuses on preserving local distances within the data. In our case, *C*_*MAN*_ = *C*_*UMAP*_ (see Equation 1). The second term penalises the repulsion between the corresponding instances in the two datasets. Because the penalty term in *Q*(*A*; *µ*) is smooth, we can use unconstrained optimisation methods to find a solution.

The penalty parameter *µ* controls the balance between the two terms. A high value for *µ* increases the importance of the correspondence information while a low value for *µ* focuses on the manifold projection of *A*. We have experienced that the penalising term can be unforgiving, so we suggest to start with a low value for *µ*. The optimal value for *µ* can also be learned using iterative methods, however optimizing *Q*(*A*; *µ*) is costly depending on the underlying manifold learning algorithm.

Currently CAML has been implemented as an extension of the UMAP algorithm because of its general applicability. The implementation is based on the model implementation of UMAP by the original author. Although CAML expands the algorithm, to our knowledge the implementation does not impose additional theoretical complexities on UMAP and does not remove performance improvement made to the algorithm. The UMAP algorithm also makes it possible to embed data based on an existing embedding. Leveraging this data transformation option together with image slicing releases the computational burden imposed by very large images.

All images are converted to the LAB color space prior to fusing. The LAB color space is a perceptually uniform color space that is often considered for microscopy imaging analysis, and it fits well with the current cost function. We have seen that this workflow provides good results and limits the computational burden on the system.

Only the dimensionality reduction step for the lymphoma MSI dataset has been done on an Intel Xeon CPU E5-2660 v2 2.20 GHz machine with 10 cores and 128 GB RAM. All other experiments have been done on a MacBook Pro with a 2,8 GHz Intel Core i7 CPU and 16 GB RAM.

### 2.4 Data measurements

For the human-lymph-node sample, cryosections of 5 m thickness were prepared and mounted on ITO glass slides. 2,5-Dihydroxybenzoic acid (2,5-DHB) was used as the matrix and applied using sublimation. The pixel size was set to 10 m, and the recorded m/z range was 620–1200 Da in positive reflector mode. The acquisition was performed with 200 lasershots/pixel and a laser repetition rate of 10 kHz, resulting in an acquisition speed of 32 pixels/s. For the mouse brain spatial transcriptomics samples, H&E images and count matrices were downloaded from https://www.spatialresearch.org/resources-published-datasets/ licensed under the Creative Commons Attribution license.

## 3 Results

With correspondence-aware manifold learning we:

i. project the high-dimensional molecular features to a low dimensional space,
ii. capture spatial correspondence between the high-resolution microscopic image and the high-dimensional molecular measurements of the same tissue sample,
iii. perform correspondence-aware manifold projection using both data modalities to obtain a fused image reflecting both modalities in one visualisation.

The general methodology is depicted in Figure 2. The high-dimensional molecular data is first reduced to three dimensions. After a registration step, this hyperspectral visualisation is used as a constraint to transform the microscopy image, resulting in a fusion of the molecular information with the microscopy data. By modeling the manifold, and piece-wise transforming the full-resolution microscopy image while taking into account the properties of the molecular data, we are able to visualise the molecular image at a much higher resolution. As such we leverage the complementarity of the high spatial resolution offered by optical microscopy with the high-dimensional but lower spatial resolution molecular imaging data.

**Figure 2:**
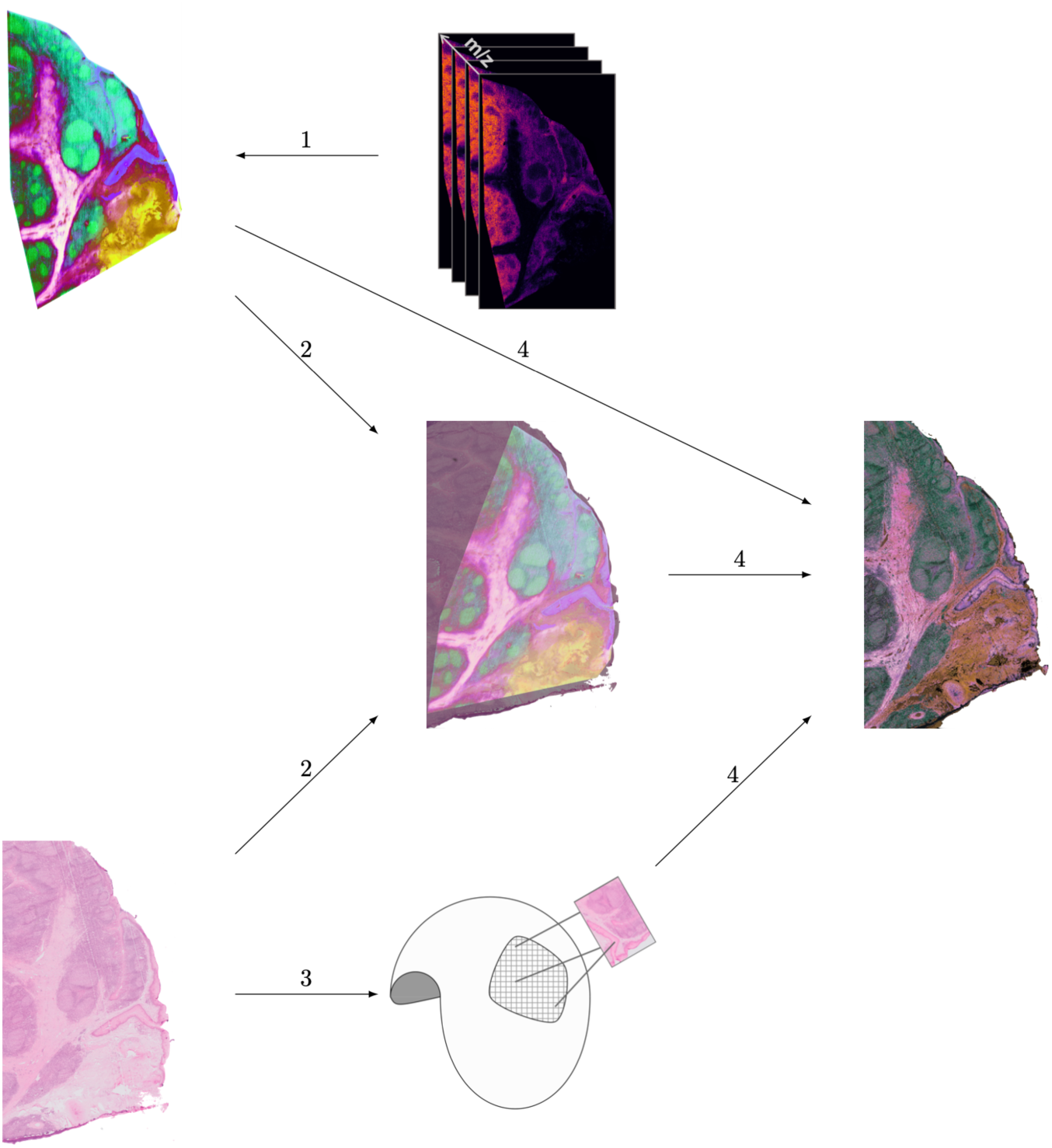
Method overview. (1) The molecular data represented by an (*n* × *p*) matrix is reduced to an (*n* × 3) target embedding. (2) The pixel coordinates of the molecular and the microscopy image undergo a registration step such that we obtain a correspondence matrix. (3) Subsequently, the microscopy image is subjected to a dimensionality reduction step, wherein each pixel is evaluated in function of its correspondence to the target embedding. (4) Specifically, the projection step in the dimensionality reduction method is constrained based on the target embedding, causing pixels in the microscopy image to receive a similar color based on the reduced target embedding of the molecular data. This approach enables the transfer of information obtained from a complete high-dimensional molecular dataset to a single microscopy slide, but can also be used to transfer the information from a single feature or molecular image to the microscopy slide.

We demonstrate the correspondence-aware manifold learning approach for data fusion of molecular measurements and their corresponding microscopy images. We show:

i. the prediction of molecular trends at a higher spatial resolution through data fusion,
ii. the prediction of molecular trends outside of the tissue area measured with out-of-sample prediction,
iii. the general applicability of the method.

### 3.1 Correspondence-aware manifold learning for data fusion of molecular and microscopy images

In Figure 3, we show the low dimensional representation of reactive lymphoid tissue in a human tonsil MSI dataset (500.000 pixels × 8000 *m*/*z* features, measured at 10*µm* resolution). This hyper-spectral visualisation represents the complete 8000 *m*/*z* feature space compressed into 3 dimensions, such that each color is connected to a molecular trend present in the data. This molecular, hyper-spectral visualisation is subsequently used to perform data fusion with the corresponding microscopy image. In Figure 4, the fused results show that the molecular trends in the data can be visualised at a much higher resolution.

**Figure 3:**
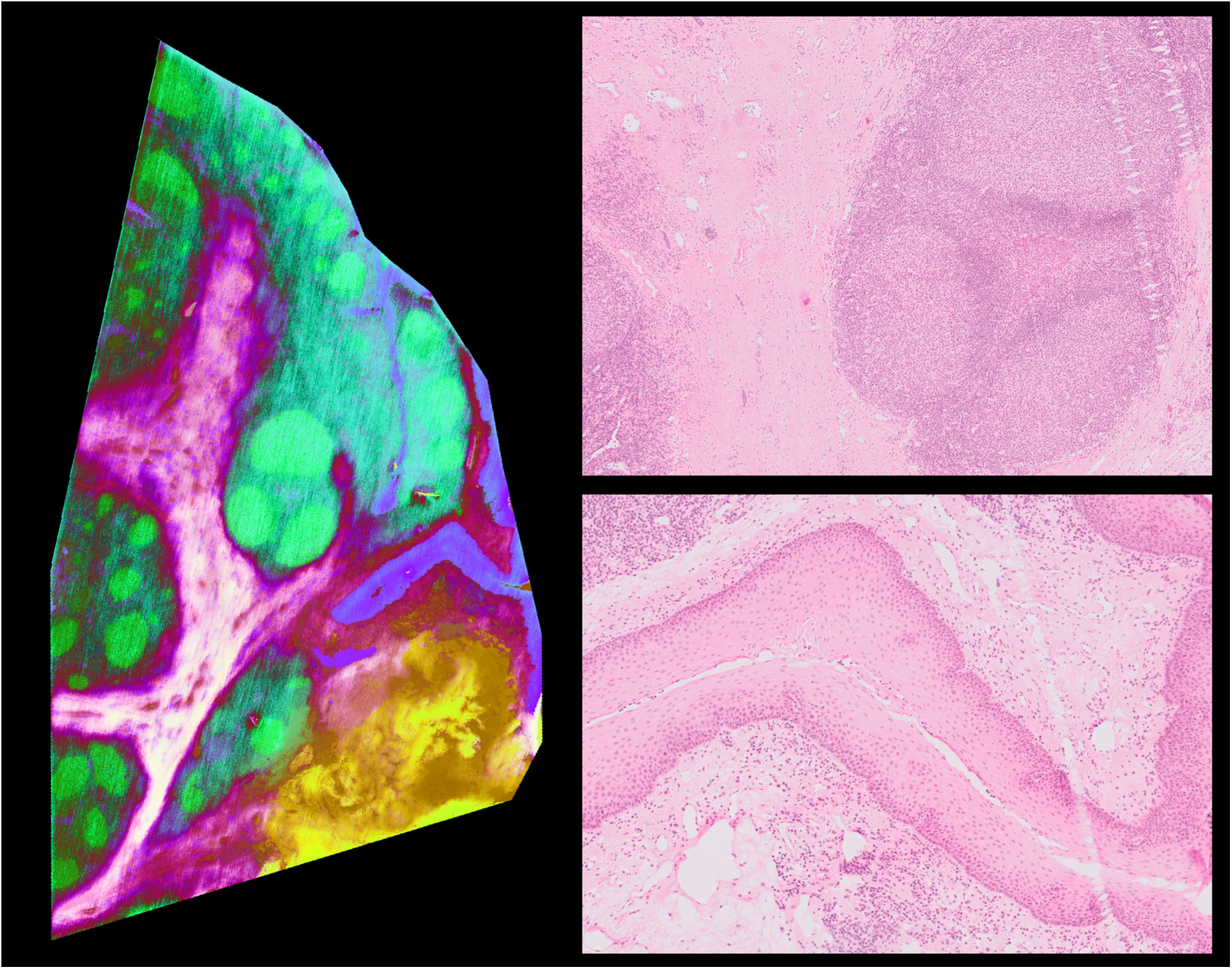
Low-dimensional representation of lymphoma MSI dataset and corresponding H&E image. (a) Shown on the left: the low dimensional representation of a human lymphoma MSI dataset (500.000 pixels × 8000 *m*/*z* features, 10*µm* resolution). The different colours reflect the molecular trends present in the data. On the right, two parts of the corresponding microscopy image are shown.

**Figure 4:**
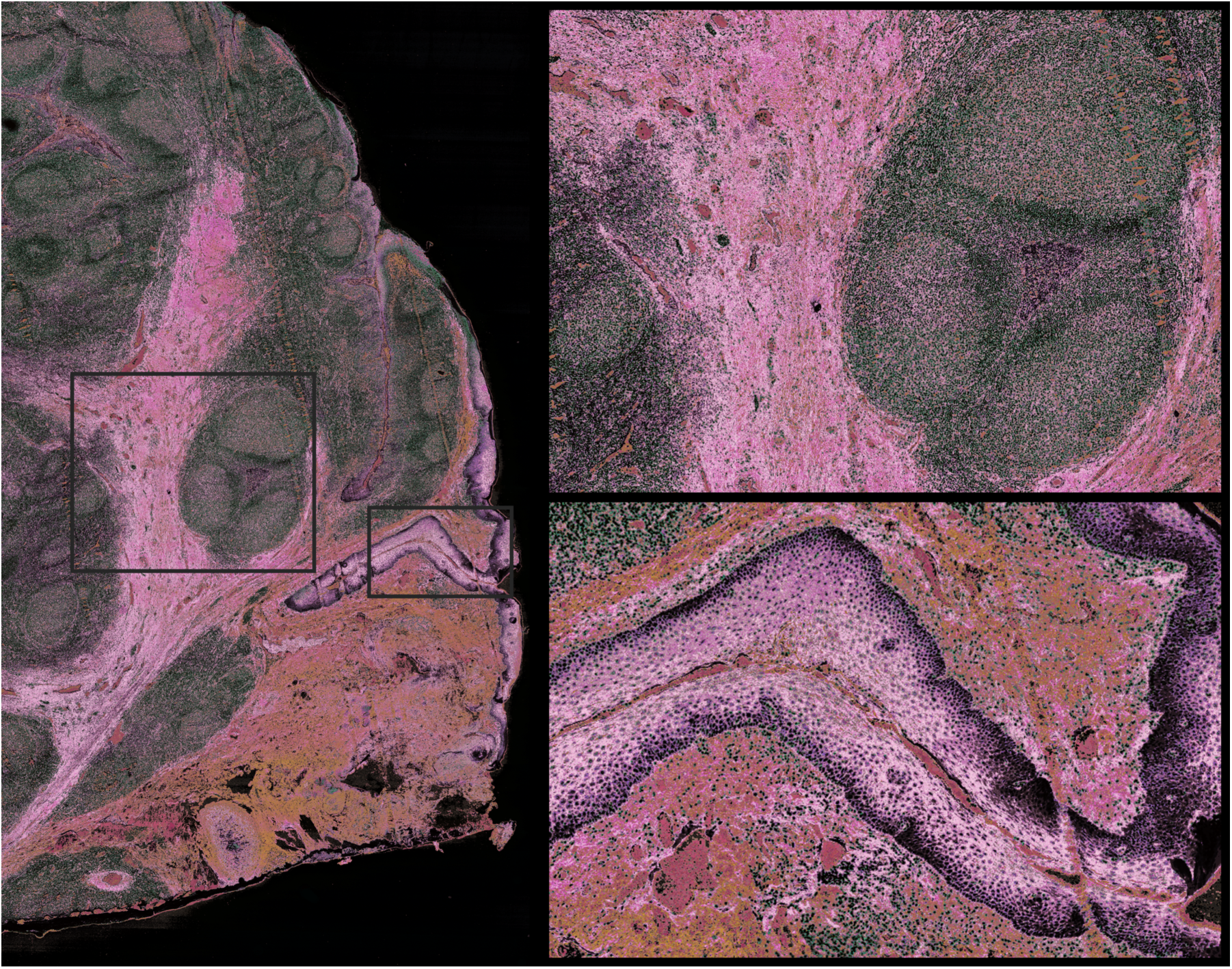
Fusion of mucosa-associated lymphoid tissue of the tonsil MSI dataset and corresponding H&E image. On the left the data fusion result for the molecular and microscopy images is shown. On the right the fusion details are shown for two regions.

The goal of performing data fusion is to exploit the complementarity present between the different data modalities such that the resulting visualisation goes beyond the information offered by a single modality. In figure 5, a fused detail is shown of the extracellular matrix and stroma comprising glycoproteins and proteoglycans with below the multi-layered squamous epithelium (panel A). Panel B shows a detailed view of a cluster of plasma cells. The nuclei of these cells received a black colour after data fusion, while the green colour seems to correspond to the cytoplasm. In active plasma cells a high density of the golgi apparatus is required for the synthesis of immunoglobulins, which explains the larger amount of cytoplasm present. A closer look reveals that these plasma cells are infiltrating the epithelium (panel C) and display a kind of integration with the present epithelial cells (purple colour) such that they do not overlay or compress these keratinocytes. Panel D shows that the keratinocyte cells further away from the basement membrane have become larger in comparison to the ones closer to this basal layer, which can be explained by the progressive maturation process taking place inside of the squamous epithelium. This maturation process is associated with changes in the composition of the cytoplasm with mainly an increase in the number of cytokeratines, which are part of the cytoskeleton. These findings are supported by the corresponding H&E stainings in panels B’-D’, and show the power of data fusion to surpass the regional or subregional level of interpretation offered via MSI by bringing the molecular information to the cellular level.

**Figure 5:**
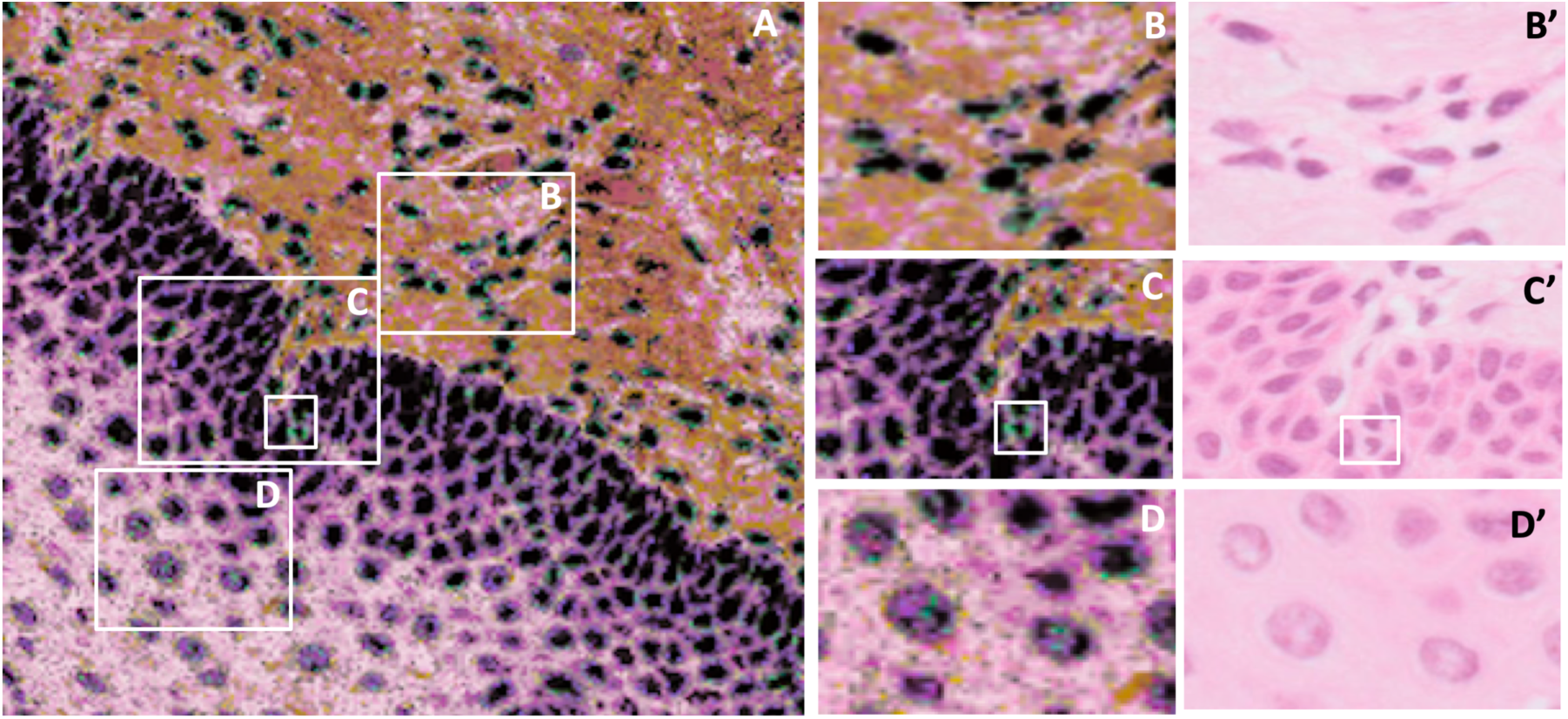
A detail of the fused data. Panel A shows the extracellular matrix and stroma comprising glycoproteins and proteoglycans, as well as plasma cells on top and the multi-layered squamous epithelium below. Panel B shows a detailed view of a cluster of plasma cells. The nuclei of these cells received a black colour after data fusion, while the green colour seems to correspond to the abundant cytoplasm. In active plasma cells a high density of the golgi apparatus is required for the synthesis of immunoglobulins, which explains the larger amount of cytoplasm present. A closer look in panel C reveals that these plasma cells are infiltrating the epithelium, as they display a kind of integration with the present keratinocytes (purple colour) such that they do not overlay or compress these cells. Panel D shows that the keratinocyte cells further away from the basement membrane have become larger in comparison to the ones closer to this basal layer, which can be explained by the progressive maturation of these cells. These findings are supported by the corresponding H&E stainings in panels B’-D’. Note that these results are provided at the cellular level: individual cells with their particular nuclei and cytoplasm are shown. This demonstrates the power of data fusion to surpass the regional or subregional level of interpretation offered via MSI by bringing the molecular information to the cellular level.

In the Supplementary figures additional examples highlight the potential of data fusion. In Supplementary figure S2, two blood vessels are shown where we can see that a venule on the left is surrounded by a thin layer of endothelial cells, while the arteriole on the right is lined by a layer of smooth muscle cells. In light pink we can also perceive some collagen fibers. Figure S1 demonstrates a secondary B-cell follicle where the reactive germinal center is surrounded by a lymphocyte corona, highlighted by an orange and green dashed line respectively. Supplementary figure S3 shows how data fusion can support us in distinguishing artefacts from true biological signals. The epithelium contains a small, rounded structure where the number of cells is increased. While this structure is visible in the H&E image, it becomes more pronounced upon data fusion.It is regarded as an artefact created during the cutting process.

All findings are supported by the corresponding H&E images. Moreover, these results highlight the potential of our method to distinguish individual cells in their micro-environment instead of being limited to the interpretation of regional or subregional molecular trends measured by MSI. This could be very valuable when evaluating for instance the invasiveness of individual tumor cells and their interaction with the tumor micro-environment.

### 3.2 Out-of-sample prediction

In addition to performing data fusion for the area covered by the MSI measurements, we can predict the molecular distributions for the entire microscopy image through out-of-sample prediction, as shown in Figure 6. This approach enables the interpretation of a much larger microscopic area based on a limited amount of molecular information. This can be a valuable asset given the high costs associated with molecular measurements or the limited amount of tissue that is often available.

**Figure 6:**
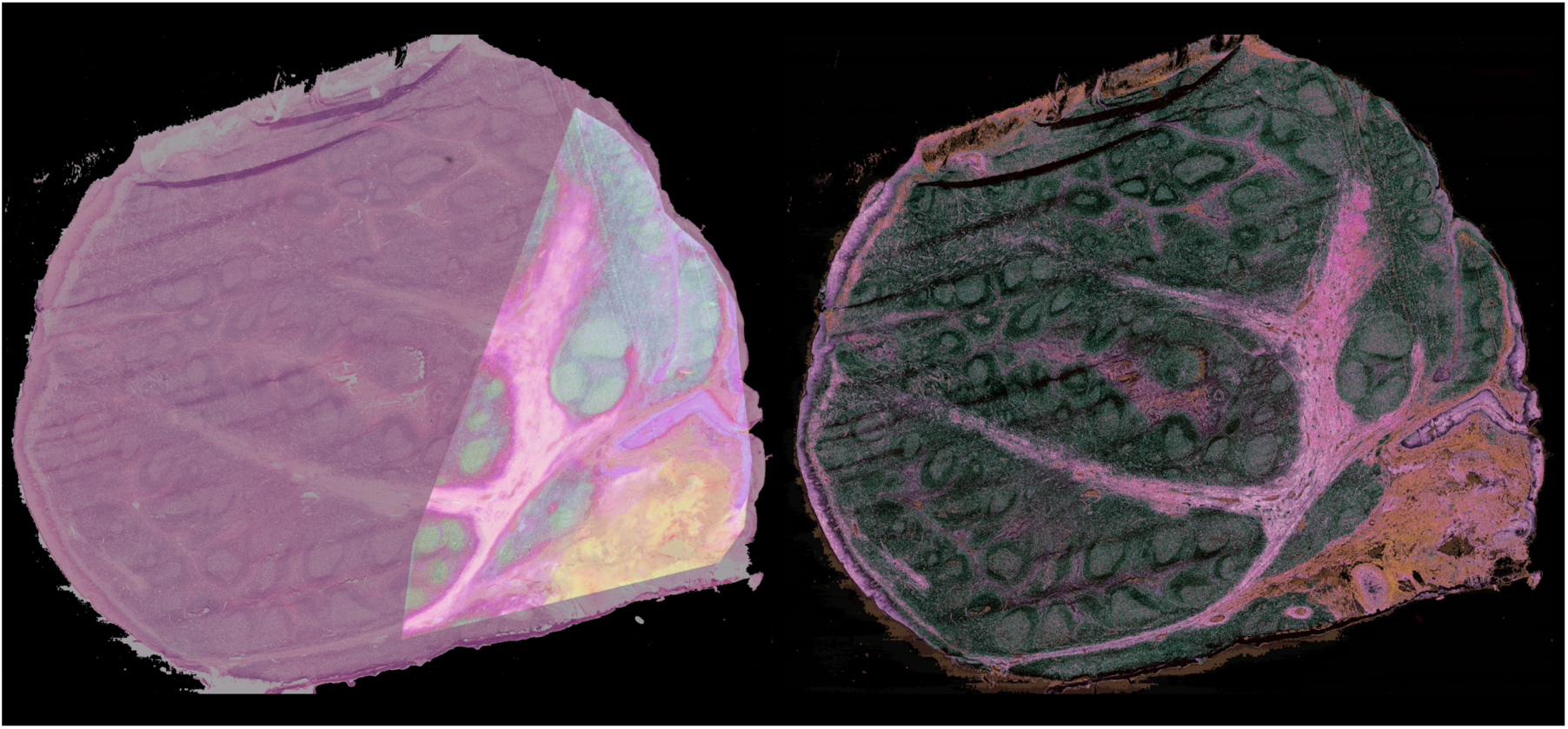
Example of Out-of-sample prediction. On the left an overlay is shown of the region in the lymphomoid tissue measured by MSI with the microscopy slide. On the right, the result of out-of-sample prediction shows the fused result for the complete microscopy or H&E image.

### 3.3 Data fusion for spatial transcriptomics data

We illustrate the general applicability of the method for spatial omics data. In supplementary figure S4, we show the hyperspectral visualisation of the low dimensional representation obtained for a spatial transcriptomics mouse brain measurement and the corresponding microscopy image with the fused result on the right. We show these results to highlight the potential of this method for other spatial omics technologies. Notwithstanding the low spatial resolution of the molecular measurement (281 pixels × 16.416 features), we can see that the green coloured cell nuclei are embedded in a purple background of the tissue center. Given the low spatial resolution, we want to emphasize the restricted potential for biological interpretation. However given the fast technical improvements that are being made in terms of spatial resolution, we believe the proposed method holds a lot of potential for data fusion of spatial omics measurements. In this light, in supplementary figure S5, we can also observe that the highlighted regions show a clear correspondence of the green dots with the cell nuclei, as stained by hematoxylin, across the associated H&E image.

## 4 Discussion

Scalable and powerful dimensionality reduction methods have become indispensable to deal with the growing number of high-dimensional datasets. Non-linear dimensionality reduction methods, such as t-SNE and UMAP, have brought and continue to bring significant value for the biomedical sciences in this regard [13, 14]. Due to their strong visualisation capabilities these methods have become a standard for the analysis of high-dimensional datasets [15]. And while the number of high-dimensional and spatial omics measurements keeps on growing, the computational methods capable of fusing the multi-modal measurements acquired from the same sample are lagging behind. In this work we present a novel data fusion method that is able to compress and fuse the complete molecular and microscopic feature spaces toward a combined image. This work builds on the framework of non-linear dimensionality reduction methods to enable the fusion of molecular and microscopic data obtained from the same tissue sample. We demonstrate our results according to the UMAP framework, but the same principle could be applied starting from similar methods such as for example t-SNE. By constraining the projection step based on the molecular target information, we are able to transform the corresponding pixels in the microscopy image accordingly, resulting in a fused representation presenting the molecular information at a much higher spatial resolution.

In Figure 4, we highlight the potential of our method for the integration of MSI data with the corresponding microscopy images. While we show that this enables us to improve the resolution of the MSI data, these colours reflect in fact an underlying group of biomolecules. As such, in previous work, we have shown that it is possible to prioritize and identify those molecules associated with a molecular trend or color [16]. This could support researchers in finding correlations between underlying biological actors and their histopathological architecture. Recent work has shown that it is possible to correlate single cell morphological features based on microscopic images with molecular information [17]. Given that the proposed method is capable of retaining the cellular morphology in the fused results, we believe it holds potential for this area of study as well. Moreover, in figure 5 we show that the proposed method can leverage MSI measurements to study tissues at the cellular level such that we can move beyond the regional or subregional insights and evaluate the presence of aberrant cells in their micro-environment. This will be of growing importance with technological advancements in terms of spatial resolution but also with the increasing demand to integrate multi-modal data measurements. In this regard, we have also included a spatial transcriptomics sample as an example of a rapidly evolving domain with a lot of potential [18]. While the molecular measurements are at the moment still of a lower resolution, it is yet possible to show that the method is widely applicable and will be able to offer more value with increasing spatial resolution. An additional advantage data fusion has to offer is the ability to better distinguish artefacts from true biological signals (supplementary figure S3. These advantages will be useful not only in the domain of molecular imaging but in general for example when dealing with other imaging technologies such as for example MRI, CT, PET, etc …).

There are several advantages of this methodology in comparison with earlier work [19]. Firstly, it is possible to take into account non-linearities and more complex interactions, which are inherent to biological systems. Secondly, it is not necessary to build a separate, predictive model for each feature (or ion image) individually. Thirdly, the data fusion result is not hindered by an imperfect multi-modal registration. This can also be seen in Figure 6. Given that registration is often a time-consuming and difficult step, this constitutes an important advantage.

To illustrate the general applicability of the method we have also performed a study on a multi-modal MNIST dataset, an extension of the well known MNIST database of handwritten digits. In supplementary figure S6 we show that we can perform data fusion on single multi-modal digits and we can also perform out-of-sample prediction based on this initial trained data fusion model.

In conclusion, data fusion facilitates the combination of complimentary data sources to obtain insights that would not be obtained from a single modality alone. Given the growing interest towards spatial multi-omics studies, this method will be valuable to enable the mapping of molecular measurements to the underlying tissue architecture at cellular resolution. Moreover, given the large costs associated with state-of-the-art molecular measurements, the out-of-sample prediction can expand the amount of information obtained from conducted experiments. Finally, due to its broad applicability we hope that the proposed paradigm will be valuable to researchers coming from different domains.

## Acknowledgements

We would like to thank Arndt Asperger from Bruker Daltonics for infrastructural support. This work was supported by KU Leuven: Research Fund (projects C16/15/059, C3/19/053, C32/16/013, C24/18/022), Industrial Research Fund (Fellowship 13-0260) and several Leuven Research and Development bilateral industrial projects, Flemish Government Agencies: FWO (EOS Project no 30468160 (SeLMA), SBO project S005319N, Infrastructure project I013218N, TBM Project T001919N; PhD Grants (SB/1SA1319N, SB/1S93918, SB/151622)), This research received funding from the Flemish Government (AI Research Program). Bart De Moor and Tina Smets are affiliated to Leuven.AI -KU Leuven institute for AI, B-3000, Leuven, Belgium. VLAIO (City of Things (COT.2018.018), PhD grants: Baekeland (HBC.20192204) and Innovation mandate (HBC.2019.2209), Industrial Projects (HBC.2018.0405)), European Commission: This project has received funding from the European Research Council (ERC) under the European Union’s Horizon 2020 research and innovation programme (grant agreement No 885682); (EU H2020-SC1-2016-2017 Grant Agreement No.727721: MIDAS); Thomas Tousseyn holds a Mandate for Fundamental and Translational Research from the ‘Stichting tegen Kanker’ (2014–083; 2019-091) and is supported by the ‘Stichting Me to You (https://www.stichtingmetoyou.be/nl/)‘

