## Supplementary figures for "Correspondence-aware manifold learning for microscopic and spatial omics imaging: a novel data fusion method bringing MSI to a cellular resolution"

Figure S1: The left panel shows a detail of the fused data with on the right the corresponding H&E image. Highlighted with a yellow dashed line are the secondary B follicles, the blue dashed line encircles the germinal center (GC) and the lymphocyte corona (LC) respectively. The nuclear-cytoplasmic ratio is higher in the LC than in the GC, which explains the denser/darker appearance of the LC. This is explained by the presence of smaller lymphoid cells, corresponding to mature, naive B-cells in the LC, compared to the larger centrocytes and centroblasts in the GC.

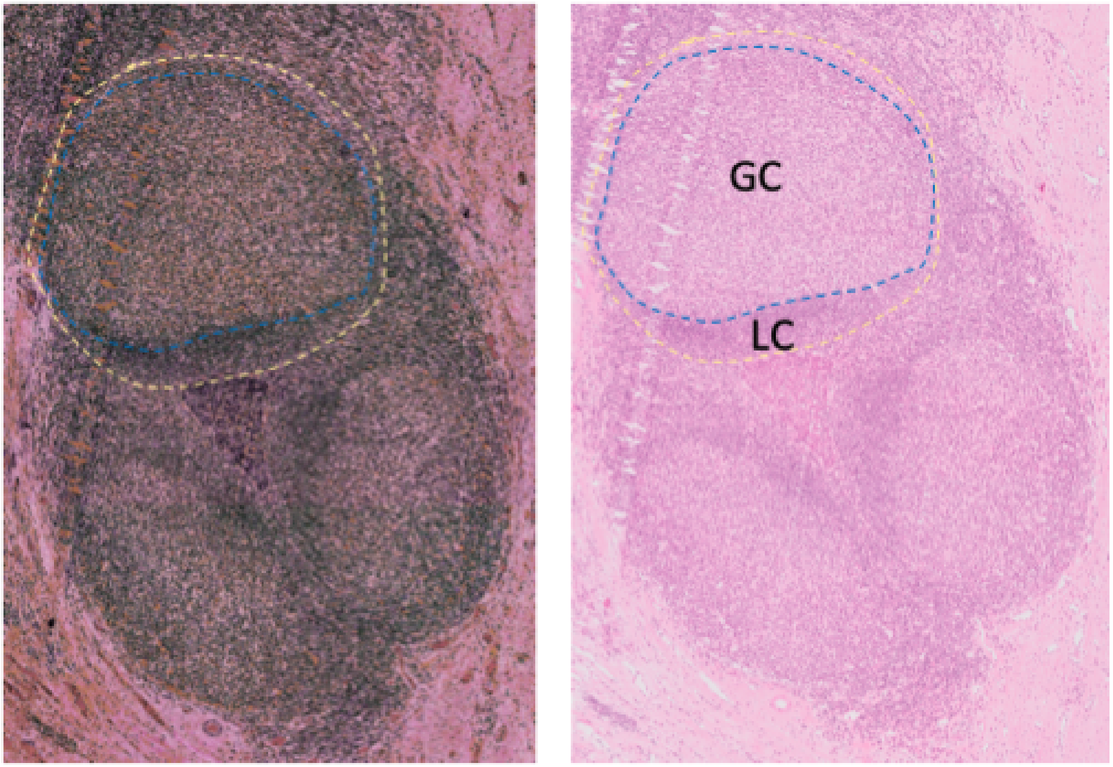

Figure S2: The left panel shows a detail of the fused data with on the right the corresponding H&E image. Highlighted in both images on the left is a venule with a thin wall that seems to correspond to a single cell layer of endothelial cells. Highlighted in both images on the right is a small arteriole with a thicker wall that also seems to contain some smooth muscle cells. The lumina of the vessels are stuffed with red blood cells. In light pink, in the stromal area between the venes and artery, we can also observe some collagen fibers.

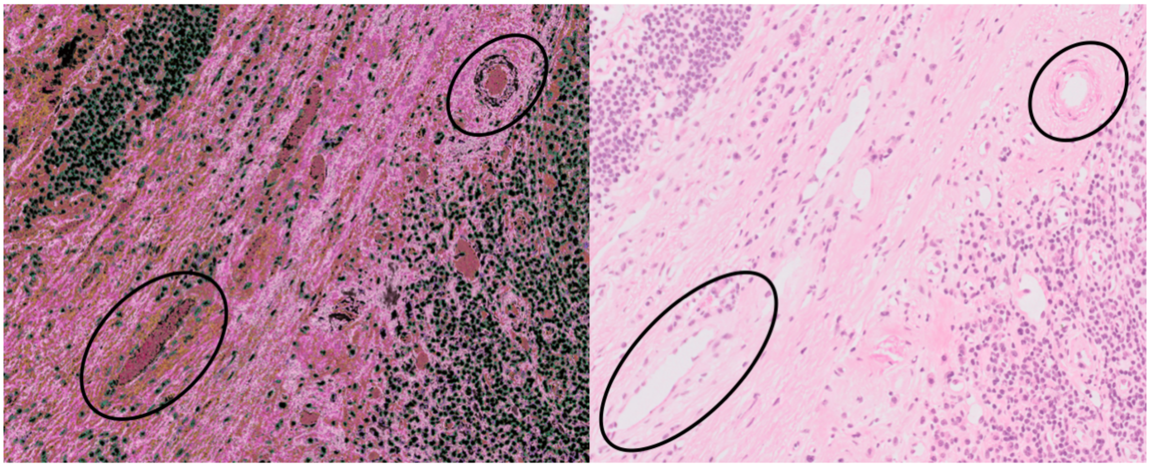

Figure S3: The left panel shows a detail of the fused data with on the right the corresponding H&E image. The epithelium contains a small, rounded structure where the intensity is increased. While this structure is also visible in the H&E image, it becomes more pronounced after data fusion. It is regarded as an artefact created during the cutting process and not a cellular structure. As such data fusion can support in distinguishing artefacts from true biological signals.

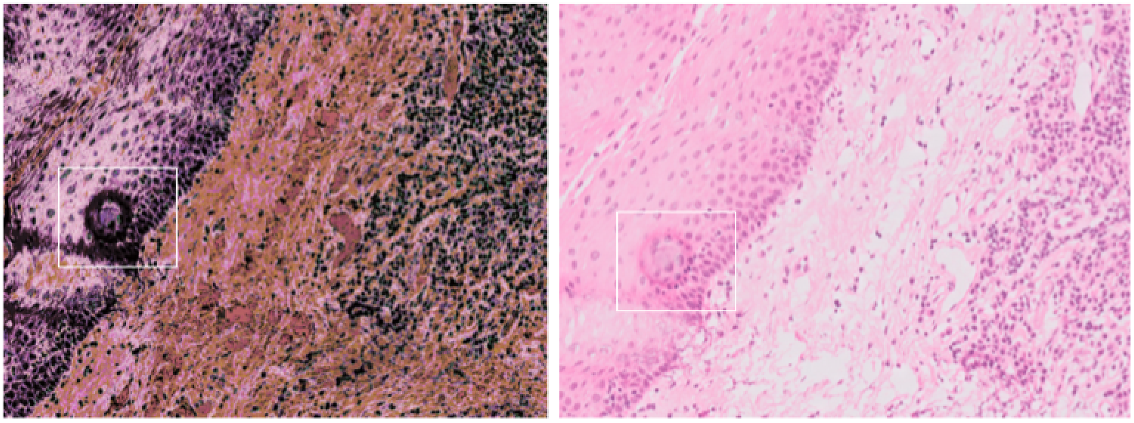

Figure S4: An example demonstrates the application of our method to a public spatial transcriptomics dataset. Shown on top:the low dimensional representation of a mouse brain (281 pixels × 16416 features, 100$\mu$ m resolution). The different colours reflect the molecular trends present in the data obtained using UMAP. Below, the corresponding microscopy image is shown, with underneath the fused result to demonstrate the general applicability. Improvements in terms of spatial resolution will make it possible to obtain results similar to the ones shown for MSI data (10 $\mu$ m resolution).

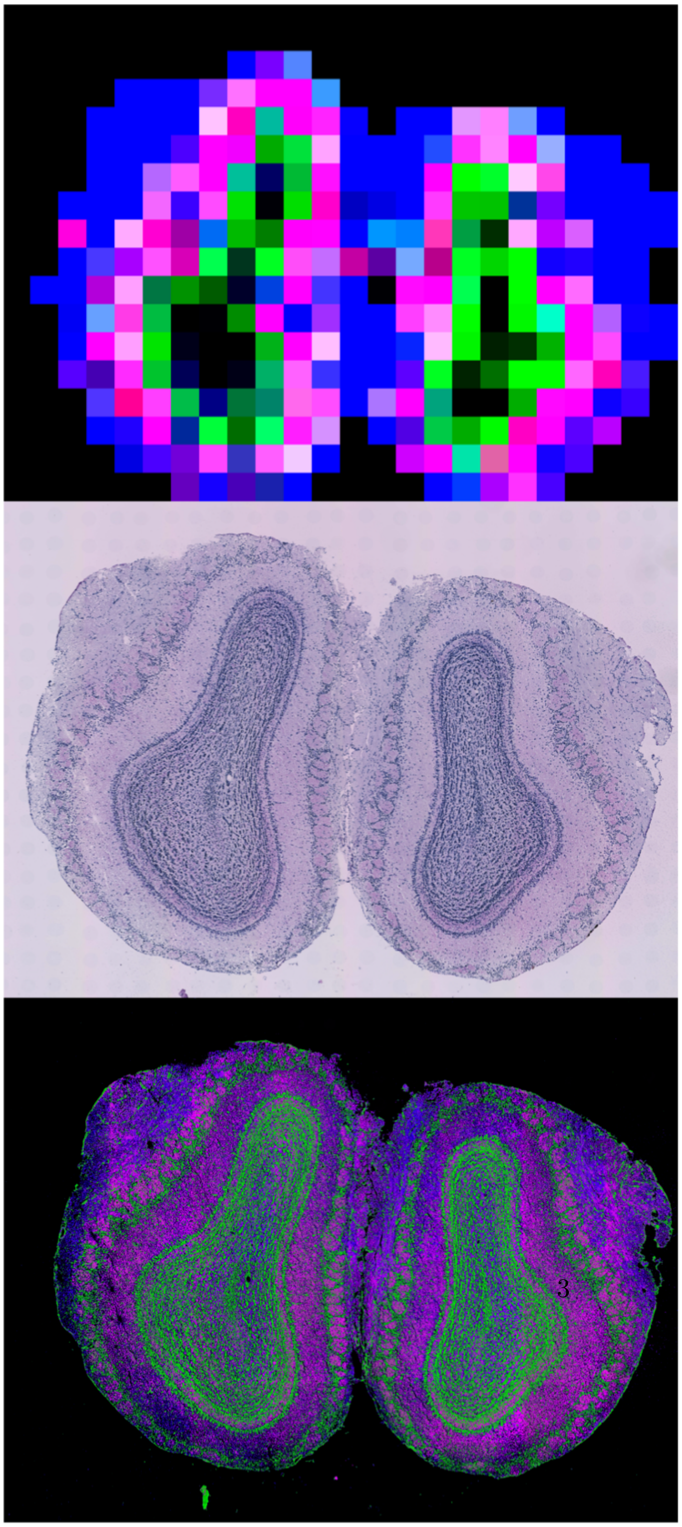

Figure S5: Illustration of consistent mapping of cell nuclei by green color upon data fusion of spatial transcriptomics mouse brain data with corresponding H&E staining. A higher spatial resolution of the data could yield more detailed results.

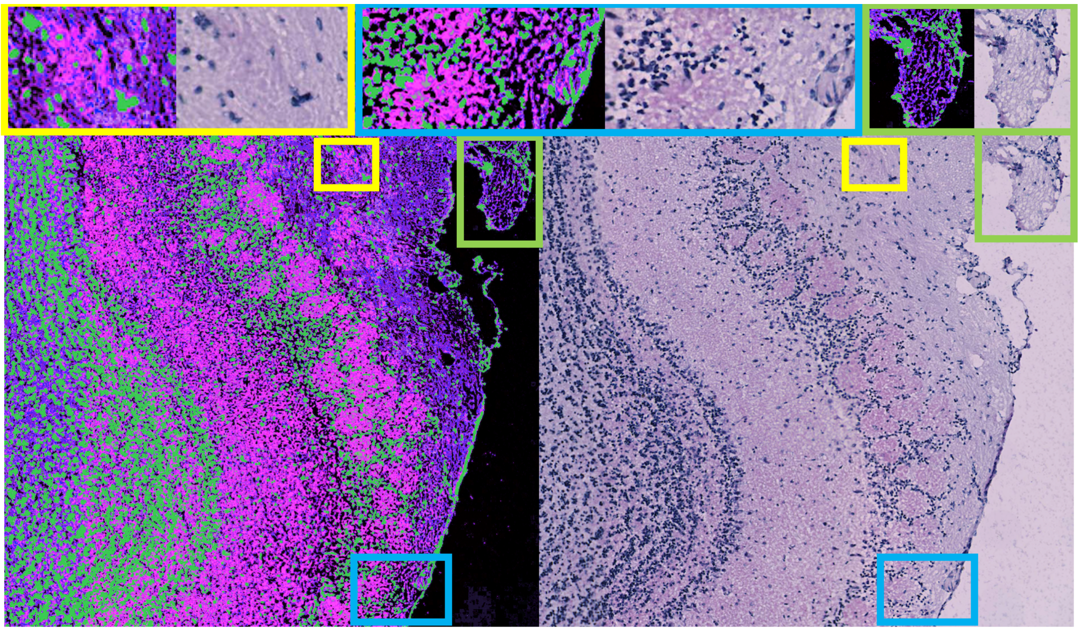

Figure S6: CAML validation and general applicability. We built a multi-modal dataset based on MNIST, the handwritten digits dataset. (1) Using a Sobel filter we created corresponding data with a shared latent space. (2) Fusion – correspondence matrix is the diagonal matrix. (3) Transform instances based on the learned model from (2). (4) UMAP embeddings of the multi-modal dataset and the fused data. We see that UMAP can still distinguish groups from the fused dataset based on the shared latent space of the two original datasets.

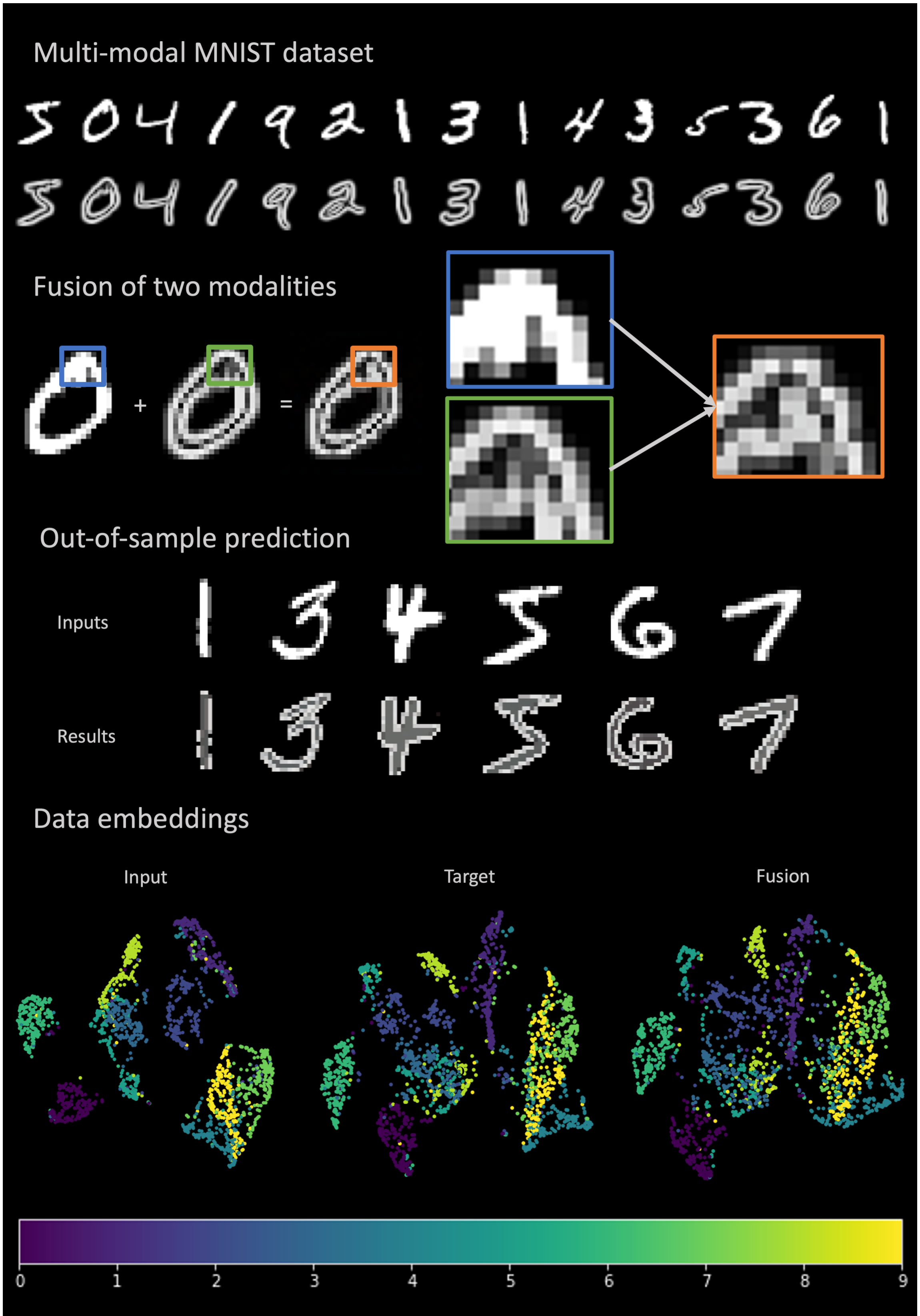
